# A novel method to select Reference Proteomes in UniProt

**DOI:** 10.64898/2026.05.12.720148

**Authors:** Pedro Raposo, Juan Sebastian Martinez Marin, Gyuri Kim, Giuseppe Insana, Dushyanth Jyothi, Jie Luo, Tanushree Tunstall, UniProt Consortium, Sandra Orchard, Martin Steinegger, Maria Martin

**Affiliations:** European Bioinformatics Institute, Wellcome Trust Genome Campus, Hinxton, Cambridge CB10 1SD, UK; Interdisciplinary Program in Bioinformatics, Seoul National University, Seoul 08826, Republic of Korea; School of Biological Sciences, Seoul National University, Seoul 08826, Republic of Korea; Swiss Institute of Bioinformatics, Centre Medical Universitaire, Geneva 4 1211, Switzerland; Department of Biochemistry and Molecular Medicine, The George Washington University, Washington, DC 20037, USA; Center for Bioinformatics and Computational Biology, University of Delaware, Newark, DE 19711, USA; Institute of Molecular Biology and Genetics, Seoul National University, Seoul 08826, Republic of Korea; Artificial Intelligence Institute, Seoul National University, Seoul 08826, Republic of Korea

## Abstract

**Motivation:** The ongoing revolution in genome sequencing is delivering an unprecedented number of genome assemblies to global repositories, resulting in an overwhelming amount of data imported to UniProt in the form of proteomes. To manage this growth sustainably, there is a need for a systematic workflow to select the best proteomes.

**Results:** We propose a novel pipeline for cellular organisms to select the best Reference Proteomes, i.e. those that best represent the protein space of a species. The pipeline uses a clustering algorithm based on MMseqs2 to select the minimum number of Reference Proteomes whilst maximising the representation of the protein space for each species. Additionally, we aligned our viral Reference Proteomes with the exemplar genome set defined by the International Committee on Taxonomy of Viruses. Because this method ensures that all species are represented with at least one Reference Proteome, the UniProt Knowledgebase increased the number of Reference Proteomes of 36% and covering 34% more species in the Tree of Life. The UniProt Knowledgebase will mainly retain proteins from Reference Proteomes and therefore this method reduces the overall number of proteins by 43%, leading to a more concise yet representative knowledgebase.

**Availability and Implementation:** https://www.uniprot.org/proteomes

**Supplementary information:** Supplementary data are available at Bioinformatics online.

## Introduction

The UniProt KnowledgeBase (UniProtKB) contains several million protein accessions from many different species across the Tree of Life, most of which are organized by their sequenced genome assembly of origin (The UniProt Consortium et al., 2024). The set of all translated protein sequences derived from a complete sequenced genome assembly is termed a proteome. Advances in sequencing technologies, together with increasing reduced costs has resulted in more genomes being submitted to public databases, exponentially increasing the number of proteomes imported into UniProt (Figure 1) (O’Cathail et al., 2024; Karsch-Mizrachi et al., 2024).

**Figure 1.**
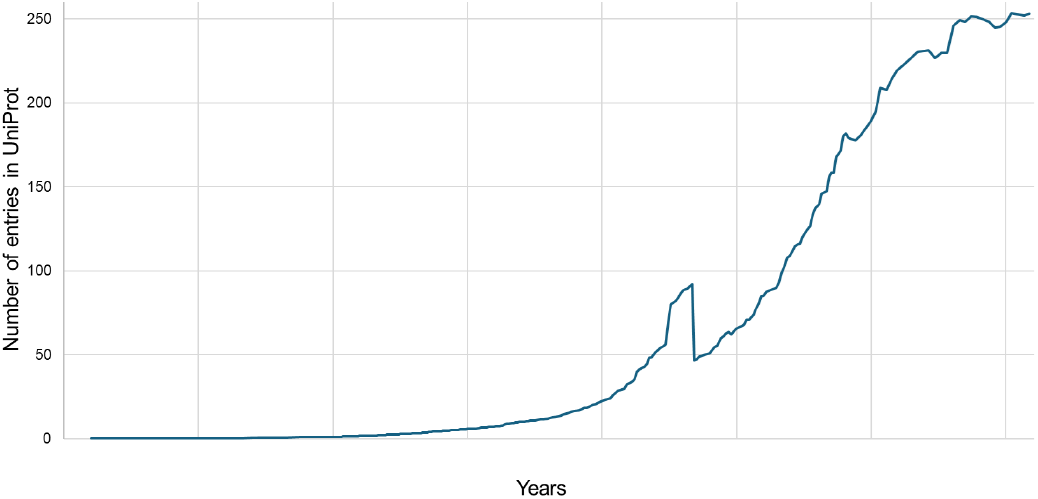
Growth of UniProtKB entries over time

This rapid growth poses long-term challenges for protein data management and leads to the publication of many redundant and poorly characterized proteomes. To prevent this, there is a need to identify, manage and annotate only the proteomes that best represent each species, i.e. Reference Proteomes (RPs). RPs are proteomes that best represent the protein sequence space for a species and/or are submitted from well-studied model organisms or are of biomedical and biotechnological research interest. By maintaining only RPs (i.e. by only retaining in UniProtKB proteins from RPs, along with some additional functionally relevant proteins), it is possible to create a sustainable dataset as its growth over the years becomes attenuated (Figure 2), and therefore UPKB will only have proteins from RPs (along with additional biologically relevant proteins), making it more concise yet representative.

**Figure 2.**
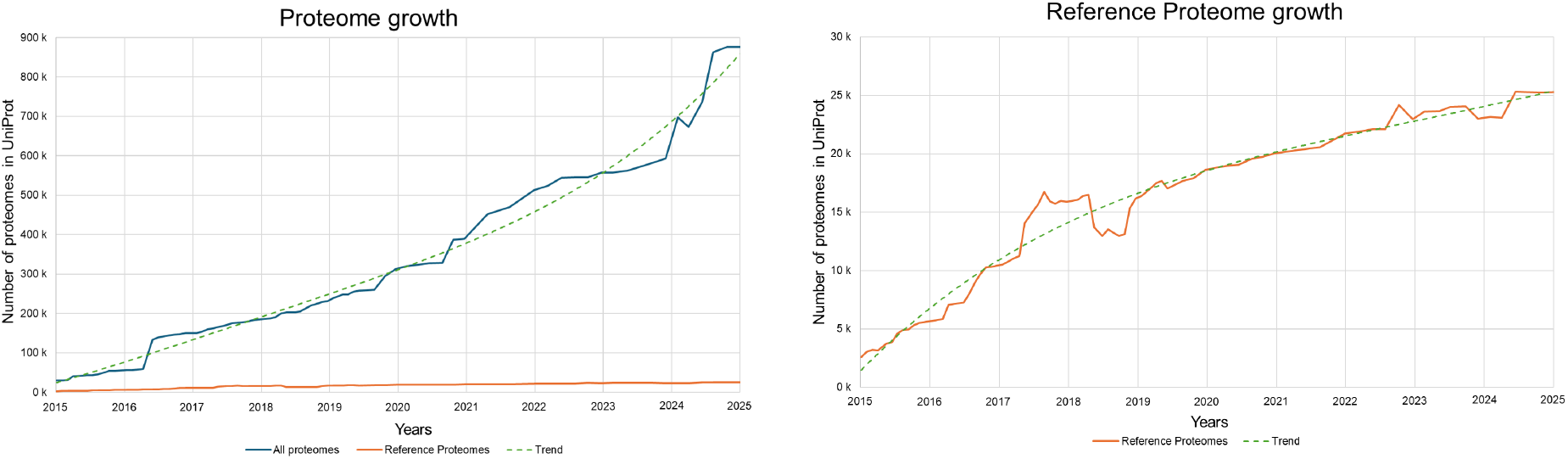
Growth trend of proteomes in UniProt throughout the years. The left plot shews corresponds to all proteomes. and it shows an exponential growth. The right plot corresponds to RPs. and it shows a logarithmic growth.

Previously, RPs were selected by a different method (Chen et al., 2011), in which it did not represent all proteome species in the Tree of Life (i.e. there were species without a single RP). This is because each RP was selected from one proteome cluster containing proteomes possibly from different species, and this resulted in some species not having a single proteome being selected as an RP, in any cluster. Also, this method does not scale well with the increasing volume of data from the International Nucleotide Sequence Database Collaboration (INSDC) (Karsch-Mizrachi et al., 2024) because the clustering step is processed by the Cluster Database at High Identity with Tolerance (CD-HIT) (Li and Godzik, 2006), in which its runtime increases exponentially with the increase of sequences being processed. To address these issues, we present a fast and scalable workflow that selects RPs for each species in the Tree of Life. This new workflow is also multi-faceted, as it replaces many previous processes built into UniProt production, such as proteome redundancy filtering and systems to prevent the import of low-quality proteomes, by achieving the same goals while simplifying the proteome production framework. The workflow incorporates previously established concepts such as proteome completeness and quality to select the best proteome(s) for each species. RP selection is automated using MMseqs2 for proteome clustering, based on protein sequences (Steinegger and Söding, 2017). Within each species, proteins from all candidate proteomes are first clustered at the sequence level, and the proteome with the highest cluster coverage is selected as the RP. The similarity between the selected RP and other proteomes is assessed to perform final clustering at the proteome level. Protein- and proteome-level clustering parameters were optimized for eukaryotic and prokaryotic taxa.

This workflow ensures the selection of at least one RP per species, and as a result, the new RP set achieves broader representation across the Tree of Life.

In addition to the analysis of cellular organisms, we present a method to identify RPs for viruses based on the International Committee on Taxonomy of Viruses (ICTV) framework (Lefkowitz et al., 2017). This approach aligns with ICTV’s responsibility for the classification and naming of virus taxa and leverages their nominated exemplar genomes, which serve as prototypic members of each viral species. Stability of RPs (and therefore UniProtKB content) between UniProt releases is crucial for maintaining integrity, reliability and traceability. While it is important to maintain data consistency from one release to the next, it is equally important to update data regularly with new information from INSDC and other genomic databases. Therefore, releases should be stable but not static, and methods were developed to address this balance.

## Methods

### 1. Proteome selection

The proteome data used for this study were the protein FASTA files of all the corresponding genome assemblies from the INSDC through the European Nucleotide Archive (ENA), as well as a set of selected proteomes from external databases such as RefSeq or Ensembl. Genome assemblies without an associated translated protein FASTA file were not included. The proteomes considered are listed in Table 1 of supplementary file 1.

The workflow considers species that meet the following criteria for analysis: (1) the taxonomic identifier must be determined to the species level, i.e. the taxonomy identifier cannot be unspecific or at the genus level (for example, “*Salmonella sp*.” corresponding to NCBI:txid 599); (2) the taxonomic identifier cannot belong to *Candidatus*).

Also, the workflow only considers proteomes that meet the following criteria: (1) its genome assembly cannot be derived from metagenomes (genomes from metagenomes were identified by either RefSeq or by UniProt internal pipelines), unless these were already RPs; (2) its genome assemblies cannot come from NCBI’s Pathogen Detection resource (https://ftp.ncbi.nlm.nih.gov/pathogen/Results/BioProject_Hierarchy/latest.bioproject_hierarchy.txt).

Quality filtering of proteomes was performed by excluding those with only a single non-nuclear genome from the analysis. Moreover, proteomes associated with genome assemblies that have been flagged by either UniProt or RefSeq quality control analyses with warnings were excluded. The exclusion reasons are listed in Table 2 of supplementary file 1.

Proteomes meeting the criteria above were clustered using MMseqs2 easy-proteomecluster to identify RPs for each species.

### 2. Parameters for MMseqs2 clustering

The following parameters were used for MMseqs2 clustering of proteomes, optimized to balance accurate protein grouping and RP selection across species. The protein sequence identity (--min-seq-id) and coverage (-c) were set as 0.9. These values were calculated from their average in protein clusters of orthologs between pairs of proteomes from the same species across the tree of life. The technical details of this analysis are shown on the supplementary file 2. The alignment coverage mode (--cov-mode), which controls the sequence length overlap, was set to 1 (“Target coverage”), to enable protein clusters to group isoforms and protein fragments with complete/canonical protein. The sequence identity mode (--seq-id-mode) was set to 0, where sequence identity is calculated solely from the alignment length without considering the length of the sequences. This ensures that sequence identity and coverage are independent variables. Clustering mode 2 (“Greedy incremental clustering mode”, --cluster-mode) was used, as it functions analogously to the CD-HIT clustering algorithm employed in the previous RP selection method. The cluster module was set to 0 (“Linclust”, --cluster-module) because it provides an efficient clustering workflow that scales linearly with input size whilst maintaining sensitivity comparable to UCLUST.

### 3. Proteome Priority Score

The Proteome Priority Score (PPS) is the parameter that determines which proteome is selected as an RP, from the proteome cluster built from MMseqs2. It was designed to consider the selection based on proteome quality through the Benchmarking Universal Single-Copy Orthologs (BUSCO) (Simão et al., 2015). Also, it keeps the changes of RPs between UniProt releases to a low amount, ensuring a stability to the UniProtKB. It is defined by four components with respective weights:

1. Manually selected Reference Proteome score: Boolean (0 or 1) indicating whether the proteome was previously manually selected as an RP by UniProt curators;
2. Representative Proteome score: Boolean (0 or 1) indicating whether the proteome was previously automatically selected as an RP by automatic methods;
3. BUSCO score: Fraction (0 to 1) indicating completeness (including single and duplication scores) inferred from BUSCO minus the fragmented percentage;
4. Representativeness score: Fraction of non-singleton protein clusters within the species (0 to 1) Clusters containing proteins from a single proteome are excluded from this calculation.

The following formula describes the PPS, with the correspondent weights:

PPS = 11,001 * Manually selected Reference Proteome score + 1,000 * Representative proteome score + 5,000 * (BUSCO complete score - BUSCO fragmented score) + 5,000 * Representativeness score

The weights are designed to prioritize certain proteomes for selection:

- The Manually selected Reference Proteomes score weight ensures that these proteomes are selected as RPs in every release;
- The Representative Proteome score weight creates a tendency for previous RPs to be selected again as RPs;
- Both BUSCO score and Representativeness score weights combined ensure the selection of a proteome as an RP if the sum of these two scores are 20% greater than a proteome that was previously marked as an RP for this species. That is, if the current proteome is 20% greater in combined BUSCO and Representativeness scores than the previously selected RP, the current proteome will become the RP. This ensures that the most complete and representative proteomes tend to be selected as RPs. The BUSCO score is split into two subscores: the complete score and a penalty score for fragments. The more fragmented the proteome, the less likely it is to be selected as RP. Regardless of BUSCO and Representativeness scores, if a proteome is a Manually selected Reference Proteome, it will continue to be an RP.

### 4. Post-process quality control checks

#### a. Taxonomy misclassification analysis

Potential taxonomic misclassified proteomes were detected. This was achieved by clustering proteomes within the same species, and also clustering proteomes within the same genus. Proteomes that share more protein clusters with proteomes from a different species (of the same genus), compared to proteomes of the same species, were identified as being potentially misclassified taxonomically.

#### b. BUSCO completeness assessment

In species where MMseqs2 selected more than one RP with significant differences in BUSCO scores, manual curation was performed. RPs with the lowest BUSCO scores in each species were evaluated and demoted to non-RP status where appropriate.

#### c. Species with abundant RPs

Manual curation was performed for species where MMseqs2 selected a large number of RPs. In these instances, literature research about the species was conducted alongside feedback from scientific community groups to guide the selection of one or a small set of RPs.

### 5. Aligning with ICTV on viral proteomes

From the GenBank assembly summary file from the NCBI FTP (https://ftp.ncbi.nlm.nih.gov/genomes/genbank/assembly_summary_genbank.txt), GenBank Assemblies (GCAs) marked as “ICTV species exemplar” were extracted, with one assembly per species. These GCAs were mapped to proteomes with translated proteins imported into UniProt. These proteomes, along with additional proteomes manually selected by UniProt curators, are designated as RPs for viruses.

## Results

### 1. Selection of non-viral RPs

From the considered 20,913 non-viral species with multiple proteomes, 224,179 proteomes were processed by MMseqs2. Species with one proteome have this one as the RP by default. MMseqs2 was run with the set of defined parameters and with the applied Proteome Priority Score on this set of proteomes, resulting in 22,491 proteomes being marked as RPs. The use of MMseqs2 has resulted in the reduction of runtime from approximately 2 weeks to 1 day, compared to the previous method.

### 2. Protein coverage from RPs

After RP(s) selection, we analyzed the distribution of the clusters being represented by the RPs, for each species. This analysis was done for proteins with different levels of frequency within the species proteomes. Among proteomes of the same species, these proteins are categorised as core (are present between 95% to 100% of the proteomes), accessory (are present between 5% and 95% of the proteomes), and rare (present between 2 proteomes and 5% of the proteomes). On average, for all domains, the RPs represent 80.55% of the core proteins, 18.87% of accessory proteins, and 0.58% of rare proteins. The high proportion of core protein clusters indicates that the majority of the species are composed of proteins that are highly conserved in all proteomes. This high core means that a small amount are accessory proteins, however the significant amount of rare proteins exists because these can be a result of sequencing and assembly artifacts, translations from genes that were horizontally transferred between species, or biological diversity between a species.

### 3. Distribution of number of RPs per species

The distribution of the number of RPs per species was investigated as the objective of the workflow is to select the least number of RPs that represent most of the proteins. From the 20,769 Eukaryotic and Prokaryotic species analysed, 19,983 (96,22%) were assigned 1 RP (15,442 (74.25%) of these are species with a single proteome, to begin with), 624 (3,00%) species were assigned 2 RPs, and 162 (0.78%) species were appointed more than 2 RPs. Ideally, this workflow selects 1 RP per species, but since there is an advantage to provide protein diversity on species, species with 2 RPs were considered to be appropriate. The species with more than 2 RPs were investigated and it was found that these are either species that were sequenced abundantly (e.g. *Escherichia coli* species has 1,629 proteomes), come from a wide range of environments (e.g. *Pseudomonas putida*), have a significant protein diversity within the same species (e.g. through bacterial gene horizontal transferring), or are parasitic or endosymbionts species which are vulnerable to genome reduction (e.g. *Buchnera aphidicola* is an intracellular symbiont of the aphid *Baizongia pistacia*). Besides these factors, small variations between proteomes might stem from artifacts from sample handling, sequencing and assembly errors and small contaminations. These species were manually reviewed by curators in order to select as few RPs as possible, to be able to represent this species.

The summary of this analysis is shown on the supplementary file 1.

### 4. Taxonomy misclassification analysis

It was determined that 248 proteomes were more similar (from the proteome cluster similarity) to a different species of the same genus, rather than with proteomes of the same species. This suggests that these proteomes are potentially misclassified: either (1) the two species genomic content is very similar (e.g. E. coli and Shigella), or (2) the taxonomy identifier provided by INSDC is incorrectly assigned to the genome. Manual curation is necessary to determine which case is true for each potential misclassified proteome. The comprehensive list of taxonomically misclassified proteomes are shown on the supplementary file 3.

### 5. Evaluation of isoform proteins clustering

The clustering system is intended to group orthologous proteins, either these being canonical and isoform sequences. To achieve this, MMseqs2 was fine-tuned on its parameters and the results were evaluated by comparing these clusters to an independent and previously established pipeline used by UniProt. This pipeline creates an automatic gene-centric mapping between entries from each eukaryotic RP that are likely to belong to the same gene. To compare MMseqs2 clusters with the gene-centric pipeline clusters, a random sample of 66 eukaryotic proteomes were chosen, and from these, their gene-centric clusters were analyzed. From these analyzed clusters, 91.95% were entirely contained within a single cluster of MMseqs2. In other words, most of the gene-centric clusters were entirely found within a single MMseqs2 cluster. This means that MMseqs2 groups isoform and canonical sequences to the same high standard as the previously established gene-centric pipeline. The summary of this analysis is shown on the supplementary file 4.

### 6. Selection of viral RPs

From the 13,053 proteomes marked as ICTV exemplar genomes by GenBank, 12,818 corresponded to proteomes with translated proteins in UniProt, where 6,836 of these were marked as new RPs. Additionally, there are 159 proteomes that were manually reviewed and marked as RPs.

### 7. Stability between releases

From release 2025_04 to 2026_01, the percentage of proteomes that got promoted or demoted as RPs (1.91%) is small compared to the percentage of proteomes that kept being RP between the releases (98.09%). This stabilizes the dataset of RPs, and is due to the fine-tuned PPS that takes account if the proteome is an RP previous to the analysis, and proteome quality (see Methods). The summary of this analysis is shown on the supplementary file 5.

## Discussion

In response to the need to keep the UniProtKB data growth stable, a new workflow for RP was developed. On cellular organisms, this was achieved by using the MMseqs2 proteome clustering workflow, and this provides many advantages: (1) The workflow allows the representation of all species in the Tree of Life, (2) it selects a minimal number of RPs, for each species, whilst representing best the protein space, (3) it is simpler and more multi-faced than previous methods, (4) it is faster and more scalable than previous methods, (5) it is traceable and contains quality control hold points for manual curation, and (6) it makes the set of RPs stable between releases but, simultaneously, not static. This workflow also promotes and maintains viral RPs, following the exemplar genomes of ICTV. All these aspects assure that UniProtKB provides to the user a good protein representation for each of these species, in a sustainable way for the future, despite the exponential growth of imported sequencing data. To ensure a biologically relevant set of RPs, each of the main parameters for the clustering were individually analyzed and carefully fine-tuned. Additionally, quality control checks are also integrated into the workflow, where the completeness and quality of the proteomes, as well as the taxonomy classification are taken into account: (1) more complete proteomes tend to become RP compared to less complete proteomes, and (2) low quality, taxonomically unclassified at the species level, or misclassified proteomes were not considered as RPs.

Starting from release 2026_02, UniProtKB contains only accessions from RPs, along with additional biologically relevant proteins. From the implementation of this workflow after release 2025_03, a 43% decrease in the number of proteins in UniProtKB is reported, despite a 36% increase in the total number of RPs. This reflects a 34% increase on species covered by at least one RP, in the Tree of Life. In summary, this work marks a major step forward in UniProt’s mission to provide the scientific community with accurate, comprehensive, sustainable, and accessible protein data, tailored to the rapidly evolving landscape of biodiversity genomics.

## UniProt Consortium members’ list

Alex Bateman, Michele Magrane, Maria Martin, Sandra Orchard, Friday Ehiguese, Aduragbemi Adesina, Emily Bowler-Barnett, David Carpentier, Paul Denny, Antonia Lock, Pedro Raposo, Conny Wing-Heng Yu, Shadab Ahmad, Chuqiao Gong, Khawaja Talal Ibrahim, Minjoon Kim, Jun Fan, Abdulrahman Hussein, Alexandr Ignatchenko, Giuseppe Insana, Rizwan Ishtiaq, Vishal Joshi, Dushyanth Jyothi, Swaathi Kandasaamy, Supun Wijerathne, Aurelien Luciani, Jie Luo, Juan Sebastian Martinez, Daniel Rice, James Stephenson, Prabhat Totoo, Vinicius de Souza, and Tanushree Tunstall at the EMBL-European Bioinformatics Institute; Alan Bridge, Paul Thomas, Lionel Breuza, Elisabeth Coudert, Ivo Pedruzzi, Sylvain Poux, Manuela Pruess, Nicole Redaschi, Lucila Aimo, Ghislaine Argoud-Puy, Andrea Auchincloss, Kristian Axelsen, Emmanuel Boutet, Cristina Casals Casas, Maria Livia Famiglietti, Marc Feuermann, Arnaud Gos, Nadine Gruaz, Chantal Hulo, Nevila Hyka-Nouspikel, Florence Jungo, Philippe Le Mercier, Patrick Masson, Sandrine Pilbout, Lucille Pourcel, Catherine Rivoire, Christian Sigrist, Shyamala Sundaram, Parit Bansal, Delphine Baratin, Teresa Batista Neto, Jerven Bolleman, Beatrice Cuche, Edouard De Castro, Elisabeth Gasteiger, Sebastien Gehant, Arnaud Kerhornou, and Monica Pozzato at the SIB Swiss Institute of Bioinformatics; Cathy Wu, Peter McGarvey, Darren Natale, Karen Ross, Yuqi Wang, Minna Lehvaslaiho, Kati Laiho, C. R. Vinayaka, Leslie Arminski, Chuming Chen, Hongzhan Huang, and Jian Zhang at the Protein Information Resource.

