## Supplementary file 2 for "A novel method to select Reference Proteomes in UniProt"

**Table of contents**

Protein identity and coverage analysis 2
Proteome similarity analysis 4
Distribution of number of RPs per species 6
Summary of changes of RPs between releases 7
References 8

### Protein identity and coverage analysis

The parameters applied on the protein cluster were analysed, for release 2025_01. Proteins that are similar in sequence should be clustered together, whilst allowing small sequence variation to be ignored. Both the protein identity and protein coverage was determined by analysing the similarity of sequences between ortholog proteins from different proteomes of the same species. A series of 187 species were randomly selected from phyla spread around the tree of life, encompassing 15 archaea, 65 eukaryota and 107 bacteria species. For each species selected, two proteomes were randomly selected. The total protein count was 3,180,617.

Searches were carried out in parallel for each couple of proteomes, in both directions (with proteome A as query proteome B as target, and also the other way around). The total number of orthologous pairs analysed were 2,421,845. The statistical measures for the comparisons are presented in Table 1. For all species, the cumulative distribution of percentage identity of best orthologous hits shows that the great majority share more than 90% sequence identity for over 90% of the orthologous hits, and this is presented in Figure 1.

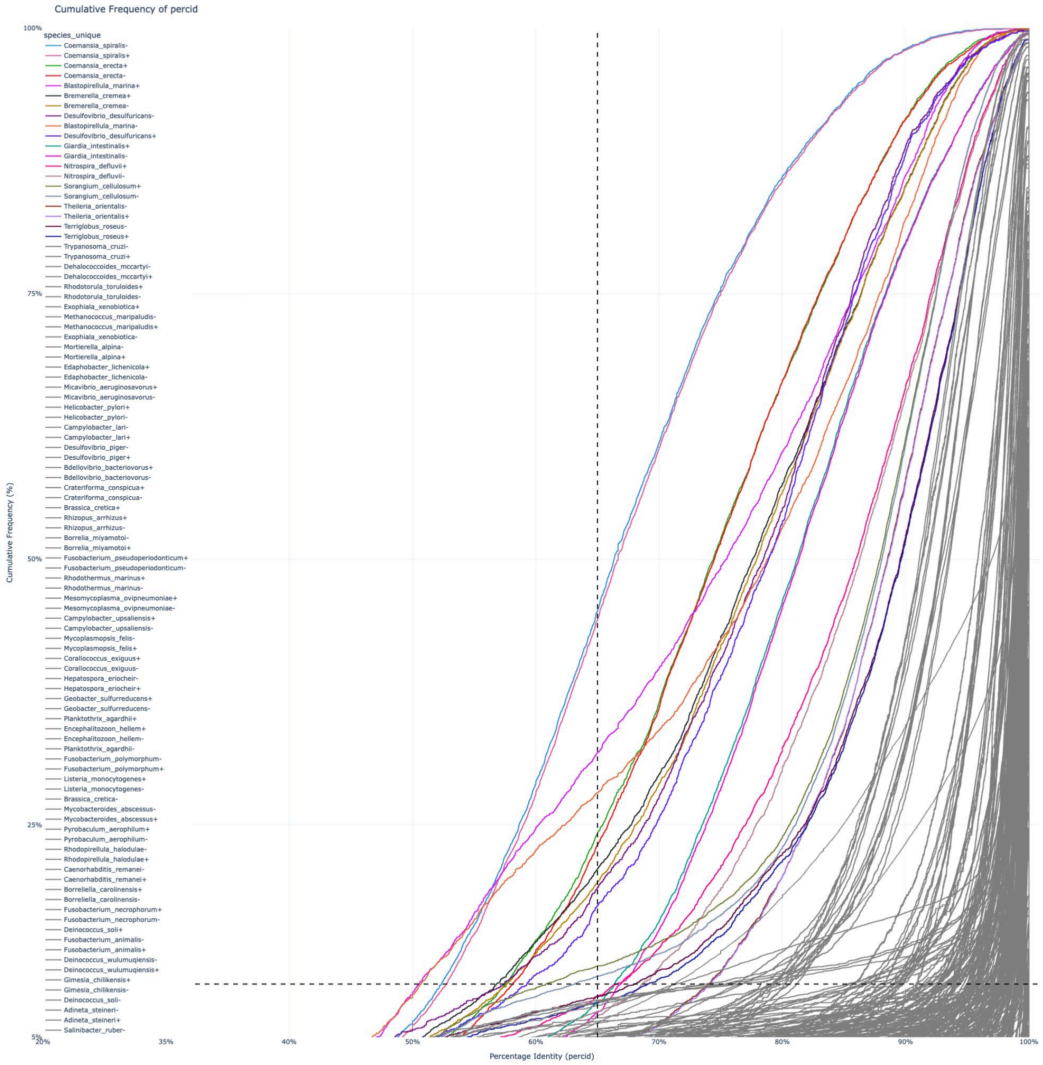


Figure 1 — Cumulative frequency plots of percentage identity by superregnum. The 10 species with lower median percentage identity are marked in color. "+" indicates forward searches (A vs B) and "-" indicates backward searches (B vs A).

With an average value of 8,551 proteins, and 6,569 orthologs between comparisons, for each species, the average sequence identity over the alignment is 96.61%, and the average coverage is 97.94%. In order to consider move variability in some species, more than others, both sequence identity and coverage were set to 90%.

The percentage identity (and alignment length as function of query and target length) for the best hits were obtained from Smith-Waterman searches of all proteins of a proteome against all proteins of another proteome. Searches were done using ssearch36 from the FASTA package with VT40 scoring matrix and E-value threshold of $1E^{-6}$.
Shallower scoring matrices like VT40 are more effective (compared with "deep" ones like BL62 and BL50) to limit the scope of the search to sequences that are likely to be orthologous between recently diverged organisms (Pearson, 2013).

Table 1 — Statistical measures of sequence identity and coverage of the comparisons of pair of proteomes.

| | Protein count | Orthologs count | percid | cov |
|---|---|---|---|---|
| Mean | 8,551 | 6,569 | 96.61 | 97.94 |
| Standard deviation | 11,494 | 7,917 | 5.11 | 2.63 |
| Minimum | 606 | 486 | 66.91 | 83.59 |
| Q1 | 2,327 | 1,959 | 96.5 | 97.54 |
| Q2 (median) | 4,152 | 3,587 | 98.11 | 98.84 |
| Q3 | 9,507 | 7,631 | 99.32 | 99.50 |
| Maximum | 68,161 | 46,150 | 100 | 100 |

percid[1] - Median sequence identity (%) of the orthologous pairs, per search
cov[1,2] - Median alignment coverage (%) of the orthologous pairs, per search

[1] - It was computed across all the best hits found in each search (each directed proteome comparison: proteome A against proteome B and vice versa, for each species)
[2] - The coverage is defined as the length of the alignment divided by the length of the smaller protein, either from the query or target proteome: alignment length/min(query sequence length,subject sequence length)

### Proteome similarity analysis

The parameters applied on the proteome cluster were analysed, for release 2026_01. There are multiple objectives to be achieved when fine-tune the proteome similarity score. This parameter represents the unidirectional similarity of the non-RP to the RP in order to be in the same proteome cluster, and as each proteome cluster has 1 RP, the number of these clusters directly determines the number of RP for each species. Firstly, the number of RP per species should be minimized. Secondly, the protein space, determined by the number of protein clusters covered by RP selected, for each species, should be maximized. Both of these variables were evaluated by the species “representativeness”, calculated as the average number of protein clusters covered per RPs, which should be maximized. Thirdly, if multiple RP are selected for the same species, the number of proteins that those RP share within a protein cluster (here called as “duplications”) should be minimized. The proteome similarity between two proteomes was calculated by using the Dice-Sørensen coefficient: $2|X \cap Y| \div (|X| + |Y|)$, where X and Y are the number of proteins from those two different proteomes.

224,179 proteomes were analyzed, from 22,491 species. 67,240 proteomes, from 6,137 species with multiple proteomes, were used to calculate the average values (within species), for the analysed variables are in Table 2. The determined proteome similarity was balanced to be 50% as the number of duplications was relatively small (10.13%), whilst covering a large portion of the protein space (90.2%). Also, the “representativeness” did not drop significantly on this proteome similarity compared to greater values, shown in Figure 2. A proteome similarity below 50% would have a lower representativeness, and a high amount of duplication.

The relative proteome similarity is the unidirectional similarity of the RP to the non-RP in order to be in the same proteome cluster, also calculated by the Dice-Sørensen coefficient. This value can be used to analyze cases when an RP clusters with non-RPs due to the former coming from a chimeric genome, and therefore having proteins of the analyzed species plus other proteins from another species. This was analyzed and as there were no instances of these in both Eukaryota and Prokaryota, so this value was set to 0%.

Table 2 — Metrics on RPs and its proteins, resulting from MMseqs2 clustering with different proteome similarity thresholds.

| | **Proteome similarity** | | | | |
|---|---|---|---|---|---|
| | **30%** | **40%** | **50%** | **60%** | **70%** |
| Number of RPs | 6,808 | 7,052 | 7,362 | 7,923 | 9,041 |
| Number of clusters covered by RPs | 26,574,641 | 26,720,806 | 26,944,995 | 27,260,906 | 27,746,008 |
| Percentage of clusters covered by RPs | 88.96% | 89.45% | 90.20% | 91.26% | 92.88% |
| Representativeness = # clusters / # RPs | 3,903 | 3,789 | 3,660 | 3440.73 | 3,068 |
| Number of duplicate proteins | 584,422 | 1,329,655 | 2,729,369 | 5,615,186 | 11,263,088 |
| Percentage of duplicate proteins | 2.20% | 4.98% | 10.13% | 20.60% | 40.59% |

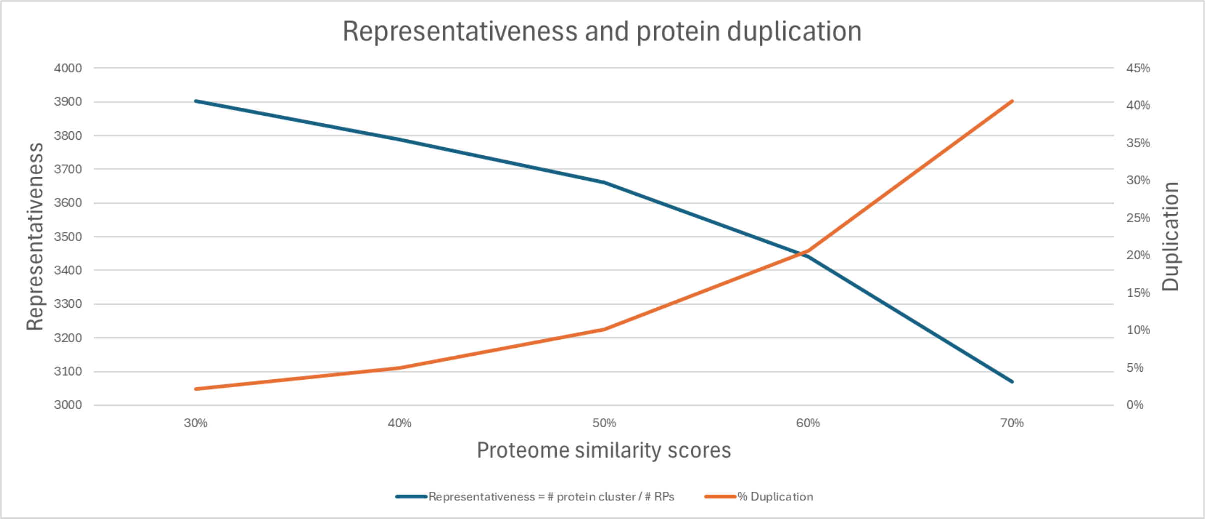


Figure 2 — Protein duplication percentage and representativeness, resulting from MMseqs2 clustering with different proteome similarity thresholds.

Figure 3 shows the average number of clusters (with at least proteins from 2 proteomes) and their population frequency, with proteome similarity of 50%. It also shows how much of these clusters are represented by the chosen RPs.

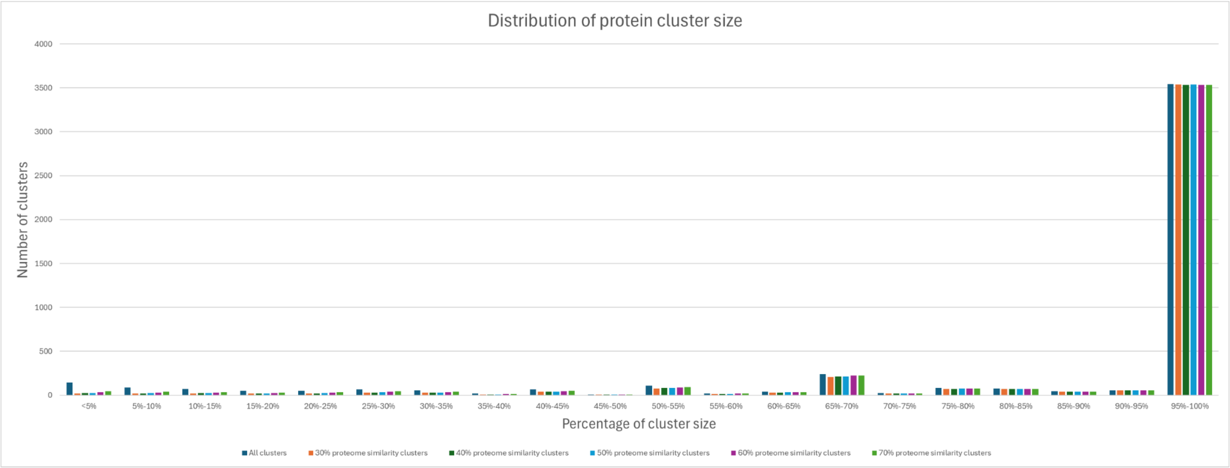


Figure 3 — Average number of MMseqs2 protein clusters, that share at least 2 proteomes, from all species of each of the Domains of the Tree of Life. This distribution is presented in a percentage of cluster size, where this size represents the number of proteins from different proteomes, i.e. the population frequency of that protein cluster. Core proteins are assigned to clusters with a population frequency between 95% and 100%, rare proteins with a frequency between 5% or less, and accessory proteins with a frequency between core and rare protein frequencies. There is a distinctive peak on the number of clusters, in the population frequency 65%-70%, as there is a large number of species with three proteomes, where there are clusters with proteins from two proteomes (population frequency of 66%). Similarly, there is a second peak in population frequency 50%-55% corresponding to species with four proteomes with clusters with proteins from two proteomes (population frequency of 50%). Dark blue columns represent all clusters, and orange columns represent the RP clusters.

All processed data from this study is found in Supplementary file 6.

### Distribution of number of RPs per species

An analysis on the abundance of RPs, per species, was done for release 2026_01.

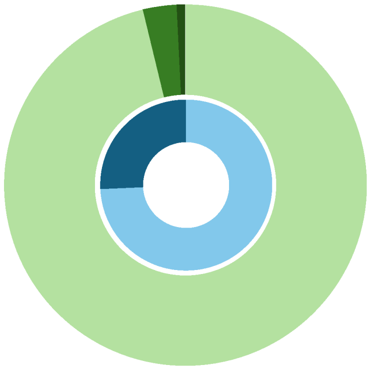

| Outer circle | Number of RPs | Number of species |
|---|---|---|
| | 1 | 19,983 (96%) |
| | 2 | 624 (3%) |
| | >2 | 162 (1%) |

| Inner circle | Number of proteomes in species | Number of species |
|---|---|---|
| | 1 | 15,422 (74%) |
| | >1 | 5,347 (26%) |

Figure 4 — Distribution of the number of RPs, per species. Most species are represented by 1 RP, some with 2 RPs, and a very small minority with more than 2 RPs.

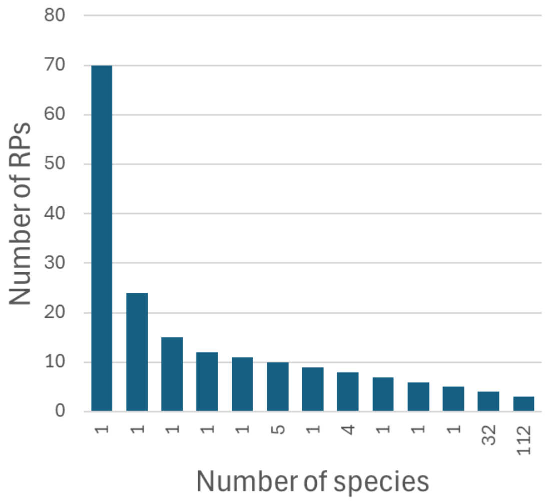


Figure 5 — Distribution of the number of species, with more than 2 RPs selected.

### Summary of changes of RPs between releases

The novel workflow was applied in release 2025_04, whereas the previous release 2025_03 contained RPs from the deprecated methods. There is an increase in the number of RPs and in the number of species with RPs, from release 2025_03 to 2025_04, in all domains.

Table 3 — Number of RPs and species with at least one RP between release 2025_03 and 2025_04, by domain, after MMseqs2 selection and manual curation.

| | **Number of RPs** | | **Number of species with at least one RP** | |
|---|---|---|---|---|
| | **Difference between releases** | | **Difference between releases** | |
| | **2025_03** | **2025_04** | **2025_03** | **2025_04** |
| Archaea | 243 | | 229 | |
| | 334 | 577 | 332 | 561 |
| Bacteria | 7,994 | | 7,518 | |
| | 9,329 | 17,323 | 9,241 | 16,759 |
| Eukaryota | 631 | | 595 | |
| | 2,767 | 3,398 | 2,671 | 3,266 |
| Viruses | 327 | | 393 | |
| | 12,817 | 13,144 | 12,584 | 12,977 |
| Total | 9,195 | | 8,735 | |
| | 25,247 | 34,442 | 24,828 | 33,563 |
